# Immune gene variation associated with chromosome-scale differences among individual zebrafish genomes

**DOI:** 10.1101/2022.06.08.495387

**Authors:** Sean C. McConnell, Kyle M. Hernandez, Jorge Andrade, Jill L.O. de Jong

## Abstract

Immune genes have evolved to maintain exceptional diversity, offering robust defense against pathogens. We performed genomic sequencing and assembly to examine immune gene variation among three zebrafish individuals. We identified remarkably high levels of sequence divergence as well as presence/absence variation among these zebrafish genomes, particularly when compared with the level of variation in human genomes. Gene pathway analysis identified zebrafish immune genes as significantly enriched among genes with evidence of positive selection. A large subset of genes was absent from analysis of coding sequences due to apparent lack of reads, prompting us to examine genes overlapping zero coverage regions (ZCRs), defined as 2kb stretches without mapped reads. Zebrafish immune genes were also identified as highly enriched within ZCRs, including over 60% of zebrafish major histocompatibility complex (MHC) genes and NOD-like receptor (NLR) genes, mediators of direct and indirect pathogen recognition. This variation was most highly concentrated throughout one arm of zebrafish chromosome 4 carrying a large cluster of NLR genes, associated with large-scale structural variation covering more than half of a vertebrate chromosome. While previous studies have shown marked variation in NLR genes between vertebrate species, our study highlights extensive variation between individuals of the same species. Our genomic assemblies also provide sequences for alternative haplotypes and distinct complements of immune genes among individual zebrafish, including the MHC Class II locus. Taken together, these findings provide evidence of immune gene variation on a scale previously unknown in other vertebrate species and raise questions about potential impact on immune function.

## INTRODUCTION

Immune genes are among the most polymorphic genes within the genomes of a large range of organisms. These genes can endow immune systems with the ability to protect organisms from rapidly changing pathogens which unpredictably attempt to evade the host response. Organisms have evolved varied complements of immune genes in order to respond effectively to these threats, harnessing a wide range of unique protein families to help ensure efficient pattern recognition by the immune system (Litman et al. 2005; Criscitiello and de Figueiredo 2013). The sequence diversity found concentrated in immune genes is often associated with positive selection and balancing selection, as populations continue to be challenged by emerging pathogens.

Adaptive and cellular immune responses are highly variable between individuals, based on extensive polymorphism, and even over time within an individual, via mechanisms such as somatic hypermutation (Flajnik 2018). A large subset of genes from the adaptive immune system specific to jawed vertebrates remain clustered within the Major Histocompatibility Complex (MHC) locus. In contrast to adaptive and cellular immune responses, innate and intracellular mechanisms to identify invaders are generally found to be more highly conserved, with many innate immune responses shared across plants and vertebrates (Turvey and Broide 2010). For example, Nod-like receptor (NLR) genes include intracellular pattern recognition molecules (PRMs) that are mediators of direct indirect pathogen recognition and other diverse functions (Maekawa et al. 2011; Meunier and Broz 2017). In the zebrafish, over 300 NLR genes have been annotated and found to be highly concentrated throughout one arm of chromosome 4 (Howe et al. 2016), making the zebrafish enriched in NLR genes compared with other vertebrates examined (Jones et al. 2016; Tørresen et al. 2018).

Representing a key model organism for developmental biology and human disease modeling, zebrafish rely on largely the same genetic pathways as other vertebrates, including humans. Zebrafish boast a high-quality reference genome, with orthologs identified for at least 80% of human disease-related genes (Howe et al. 2013). However, unlike other model organisms such as inbred mice, laboratory zebrafish have generally been maintained as outbred populations, with repeated introduction of fish from wild and captive-bred populations to help maximize genetic diversity (Brown et al. 2012; Butler et al. 2015), likely impacting diversity of immune genes.

Previously we described divergent haplotypes of the zebrafish core MHC locus, where paralogs throughout the antigen processing pathway have been maintained via balancing selection for half a billion years on alternative haplotypes (McConnell et al. 2016). This included alternate sets of immunoproteasome subunit genes and transporter associated with antigen processing (TAP) genes, as well as Class I MHC genes for antigen presentation. MHC genes in zebrafish had previously been shown to have evolved into distinct complements of genes that varied markedly between individuals (McConnell et al. 2014). Finding similar extensive variation across the linked immunoproteasome and TAP genes was unanticipated and at a scale that was unlike previous species that had been examined. Building on our previous work that uncovered significant variation in zebrafish Class I MHC pathway genes, the goal of this study was to survey additional genetic diversity in immune genes and throughout the zebrafish genome.

## RESULTS

To examine immune gene variation in the zebrafish genome and compare with variation in the human genome, we performed deep (50-60x coverage) whole genome sequencing for two clonal zebrafish lines, CG1 and CG2, in addition to a third partially inbred individual, AB3, all derived from the AB genetic background (http://zfin.org/ZDB-GENO-960809-7). Approximately 11 million single nucleotide variants (SNVs), and 2 million small insertions or deletions (indels) were called per individual using GATK HaplotypeCaller (Supplementary Figure 1). These raw variants were then hard filtered (Supplementary Table 1) to enrich for high confidence variants, yielding 6.3-7 million SNVs per zebrafish individual, substantially more than the number of SNVs (2.4-4.2 million) found in each of three different human samples (Supplementary Table 2). When adjusted for genome size, the zebrafish samples had higher SNV density, at 4.7-5.2 SNVs per kb, compared with the SNV density of 1.0-1.7 SNVs per kb in each human sample.

As expected, for both SNVs (Fig 1A) and indels (Fig 1B), the vast majority (96-97%) of filtered variants were called as homozygous for the two clonal zebrafish lines (Supplementary Table 3). Similarly, most variants were also called as homozygous (99% of SNVs and 92% of indels) for the haploid human hydatidiform mole sample CHM1, consistent with prior studies (Zook et al.2014). In contrast, filtered variants were more often called as heterozygous for the partially inbred AB3 zebrafish (56% of SNVs and 51% of indels), as well as for the two human samples of European and African ancestry (60-66% of SNVs and 56-68% of indels).

**Figure 1.**
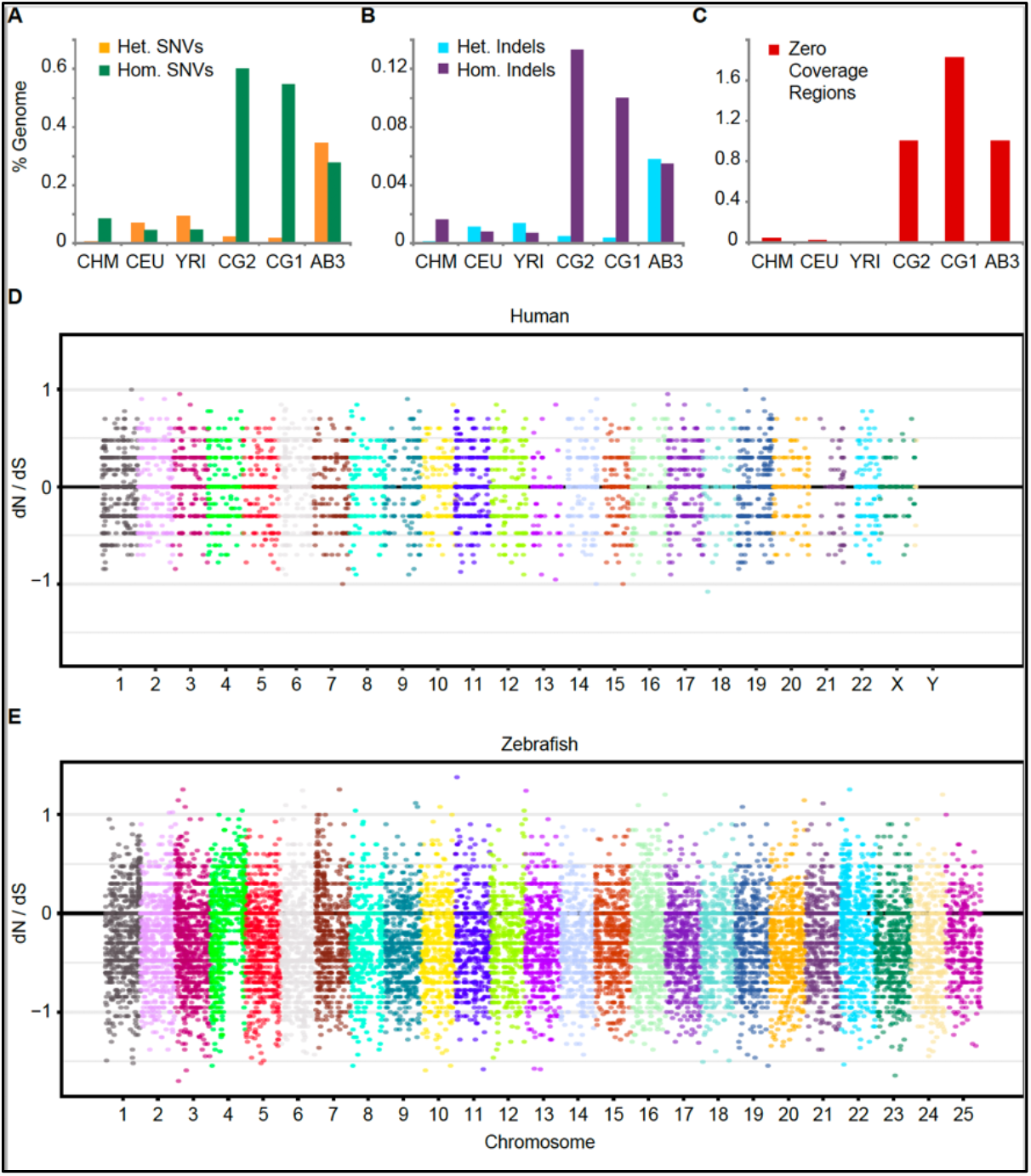
Sequence variants and evidence of positive selection. *(A)* Single nucleotide variants (SNVs) or *(B)* small insertions/deletions (Indels) were identified using GATK haplotype caller, and reported as a percentage of each genome. Both heterozygous (Het.) and homozygous (Hom.) variants are shown. *(C)* Percentage of base pairs in each genome covered by Zero Coverage Regions (ZCRs), defined as no reads mapped over ≥ 2 kb intervals. Manhattan plots of the ratio of non-synonymous to synonymous mutations (dN/dS) per allele for three human *(D)* or zebrafish *(E)* individuals. Each dot represents one gene with variants. The black horizontal line at ‘0’ indicates alleles under neutral selection, i.e. those having a dN/dS ratio of 1 (the ratio for each allele is plotted on a log10 scale). A large fraction of genes throughout the right arm of zebrafish chromosome 4 have evidence of positive selection (dN/dS > 1). CHM, CEU, and YRI are samples from the 1000 genomes project representing: a haploid complete hydatidiform mole, CHM1; Utah Resident (CEPH) with European Ancestry, NA12878; Yoruba in Ibadan, Nigeria, 19240; respectively. CG2 and CG1 are clonal zebrafish lines, and AB3 is a partially inbred fish, all on the AB genetic background.

To examine potential selection pressure, we annotated filtered variants using ENSEMBL’s Variant Effect Predictor (VEP) v85 (McLaren et al. 2016). Non-synonymous (dN) and synonymous (dS) SNVs were counted per gene, across each allele among the three human and three zebrafish samples. Variants were identified in a total of 11,201 human and 19,520 zebrafish genes. Of these, 8,544 human and 18,612 zebrafish genes had synonymous variants (dS >0), with an average of 1.4 non-synonymous and 1.6 synonymous variants per human gene, and 4.0 non-synonymous and 7.6 synonymous variants per zebrafish gene.

Evidence of positive selection was inferred for genes with a composite dN/dS ratio greater than 1 (Fig 1D, E). Not only were more variants identified per zebrafish gene, but more zebrafish genes than human genes had evidence of positive selection (3013 to 1568 genes). A total of 30 zebrafish genes had dN/dS ratios ≥ 10, compared with only two human genes. These results provide evidence that some of the high variability within the zebrafish genome has evolved under positive selection.

Similar to SNVs, relatively high numbers of small insertions and deletions (indels) were found in the zebrafish genomes. The number of indels that passed our conservative filters exceeded 1.5-2 million per zebrafish genome, in a genome size of 1.4 Gb, for a rate of approximately 1.1-1.4 indels per kb (Fig 1B). By comparison, the number of indels identified per human genome was 0.5-0.7 million (The 1000 Genomes Project Consortium 2015), in a genome size of approximately 3 Gb, for a rate of approximately 0.2-0.3 indels per kb (Supplementary Table 4). Thus, both indels and SNVs were ∼3-5-fold more abundant in the zebrafish genomes compared with the human samples analyzed.

Despite high variant density found throughout the zebrafish genome, a surprisingly large number of zebrafish genes had no variants called. Manual inspection of their sequences revealed that many of these genes lacked high quality mapped reads. This lack of aligned reads could be due to these sequences being absent from an individual zebrafish genome, or due to high divergence of the sequences relative to the reference genome. To help identify affected genes potentially overlooked by the variant calling pipeline, we looked for 2kb or larger gaps without any mapped reads, representing zero coverage regions (ZCRs) previously associated with structural variation (Pucker et al. 2016). A substantially larger fraction (>10 fold higher) of each zebrafish genome was covered in ZCRs compared with human genomes (Fig 1C, Fig 2). The relative prevalence of ZCRs offers evidence of additional variation in the zebrafish genome, highlighted by unexpectedly large stretches of reference sequence with no aligned reads.

**Figure 2.**
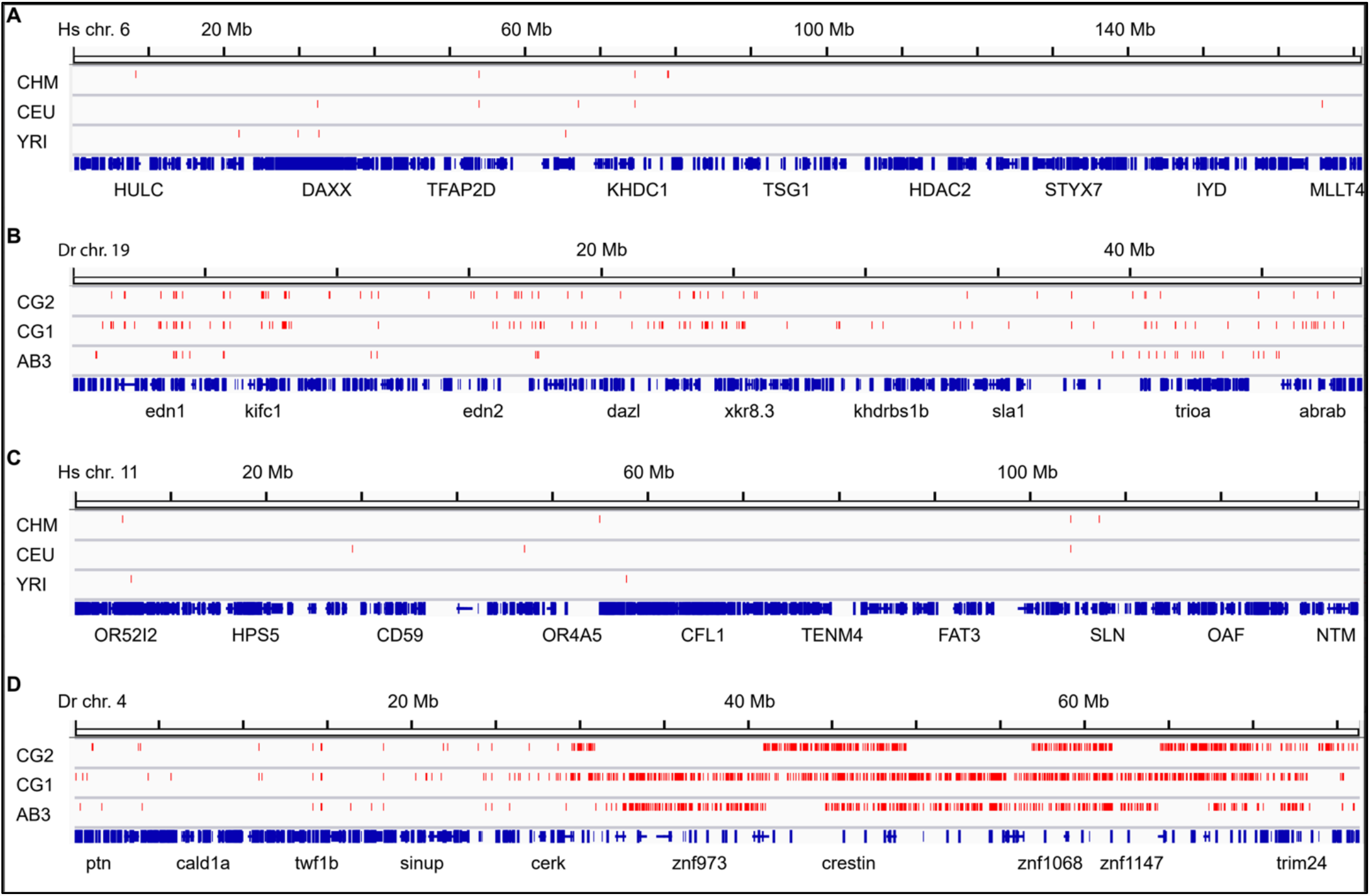
Chromosomal distribution of Zero Coverage Regions. Comparison of (A) human chromosome 6 (location of the human MHC locus), (B) zebrafish chromosome 19 (location of the zebrafish core MHC locus), (C) human chromosome 11 (location of 4 out of 25 human NLR gene family members), and (D) zebrafish chromosome 4 (location of over 300 zebrafish NLR genes). Zero Coverage Regions (no mapped reads over ≥ 2 kb intervals) are displayed in red. Gene annotation is shown in blue with a small number of genes labeled. ZCRs are found more densely in zebrafish chromosomes compared with human chromosomes and a large concentration of ZCRs is distributed throughout the heterochromatic right arm of zebrafish chromosome 4 with evidence of haplotypic differences between individuals.

Most striking was our observation of very dense ZCRs in the late-replicating, heterochromatic arm of zebrafish chromosome 4 (Fig 2D). This arm of zebrafish chromosome 4 has been a focus of annotation efforts that revealed a large number of immune genes, including hundreds of NLR genes (Howe et al. 2016). Here we find that a large percentage of these NLR genes lacked mapped reads across broad expanses, in patterns that were largely unique to each sample, indicating highly divergent haplotypes between individuals. Interestingly, this arm of chromosome 4 has been identified as linked to a sex determination region that was lost during the domestication of zebrafish (Wilson et al. 2014), suggesting different selection pressure on this portion of the laboratory zebrafish genome. This region of zebrafish chromosome 4 is highly enriched for genes with evidence of positive selection (Fig 1E).

To identify patterns of selection genome-wide, we performed gene pathway enrichment analysis via Gene Ontology (GO) annotation using ClusterProfiler (Yu et al. 2012), based on combined lists of genes with evidence of positive selection (dN/dS > 1), or on combined lists of genes with coding regions overlapped by ZCRs. These results were summarized using ReviGo (Supek et al. 2011), which revealed that immune gene pathways including antigen processing and presentation were highly enriched among the zebrafish genes with evidence of positive selection (Fig 3A), as well as with genes with exons overlapping ZCRs (Fig 3B) (supplementary Excel file). In contrast, human genes associated with positive selection (Fig 3C) or ZCRs (Fig 3D) were significantly enriched in genes involving sensory perception, or keratinization, respectively. Many of the human genes overlapping with ZCRs in our study, such as *LCE3B* and *LCE3C*, are associated with antimicrobial activity and implicated in psoriasis and wound-healing, represent variants with known presence/absence variation among human populations (The Wellcome Trust Case Control Consortium et al. 2010).

**Figure 3.**
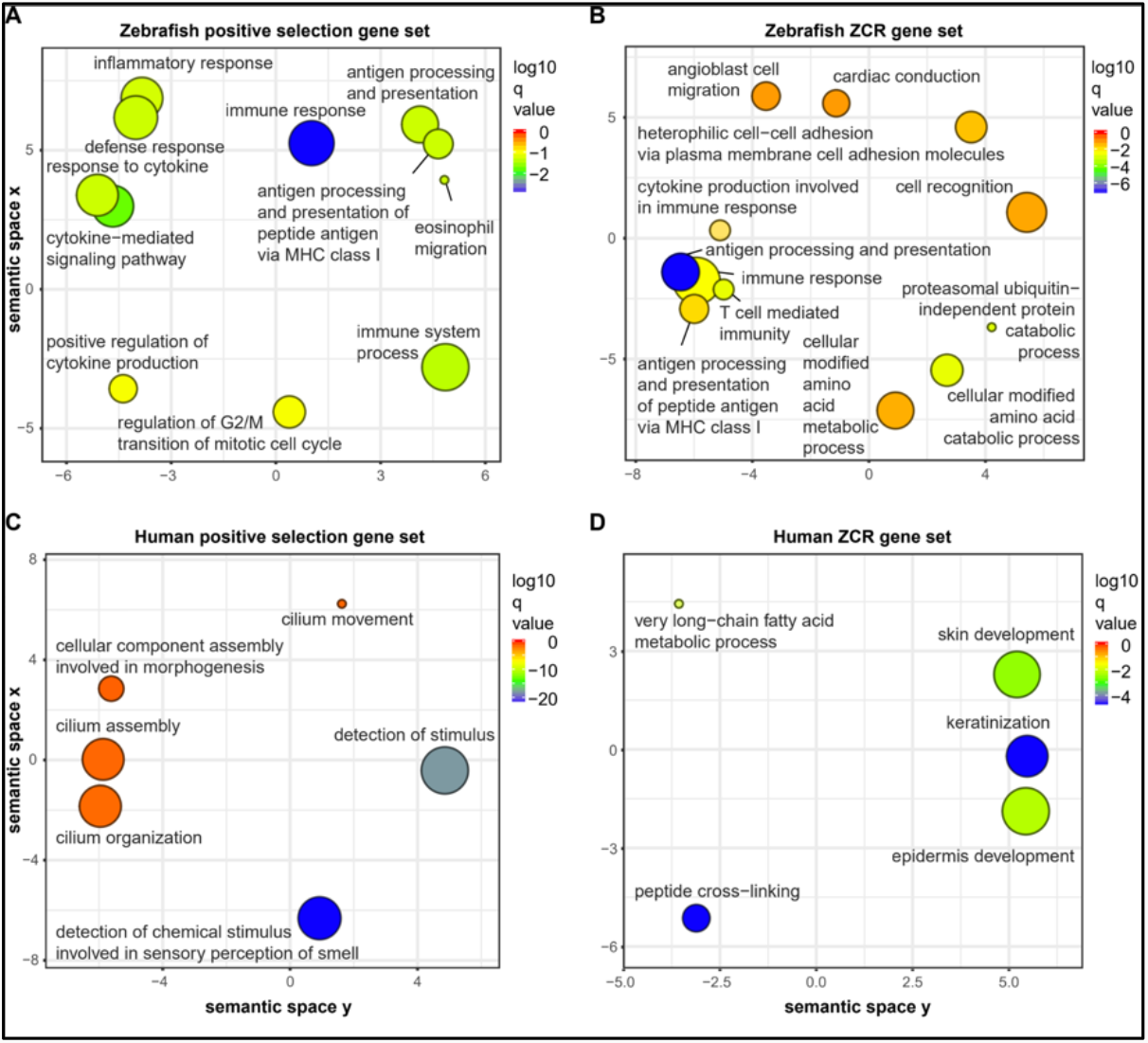
Gene annotation enrichment analysis. Genes with evidence of positive selection (dN/dS > 1) or genes with at least one exon overlapping Zero Coverage Regions (ZCRs, without any mapped reads over ≥ 2 kb intervals) were analyzed using GO (Gene Ontology) annotation to identify genes enriched in specific biological processes. Data are shown for zebrafish genes under positive selection (A), zebrafish genes overlapping ZCRs (B), human genes under positive selection (C), and human genes overlapping ZCRs (D). Gene lists are available in supplementary Excel file.

In many cases manual inspection of zebrafish and human genes found enriched in ZCRs revealed large, continuous regions of missing coverage, with clear boundaries. In other cases, coverage of mapped reads was more sporadic, with ZCRs unable to capture the smaller regions of missing coverage. Despite intermittent stretches of low or no coverage, exons for some human genes were narrowly missed by ZCRs, for example, MHC Class II gene *HLA-DRB5* (Fig 4A), a gene known to have presence/absence variation. Other human genes such as *LCE3B* and *LCE3C* (Fig 4B) overlapped ZCRs in some samples, for example, a 30kb nearly continuous region linked to psoriasis that was identified in the CHM1 haploid genome. Our diploid human samples often had relatively low coverage over these same regions, consistent with being heterozygous for alternative haplotypes lacking the reference sequences.

**Figure 4.**
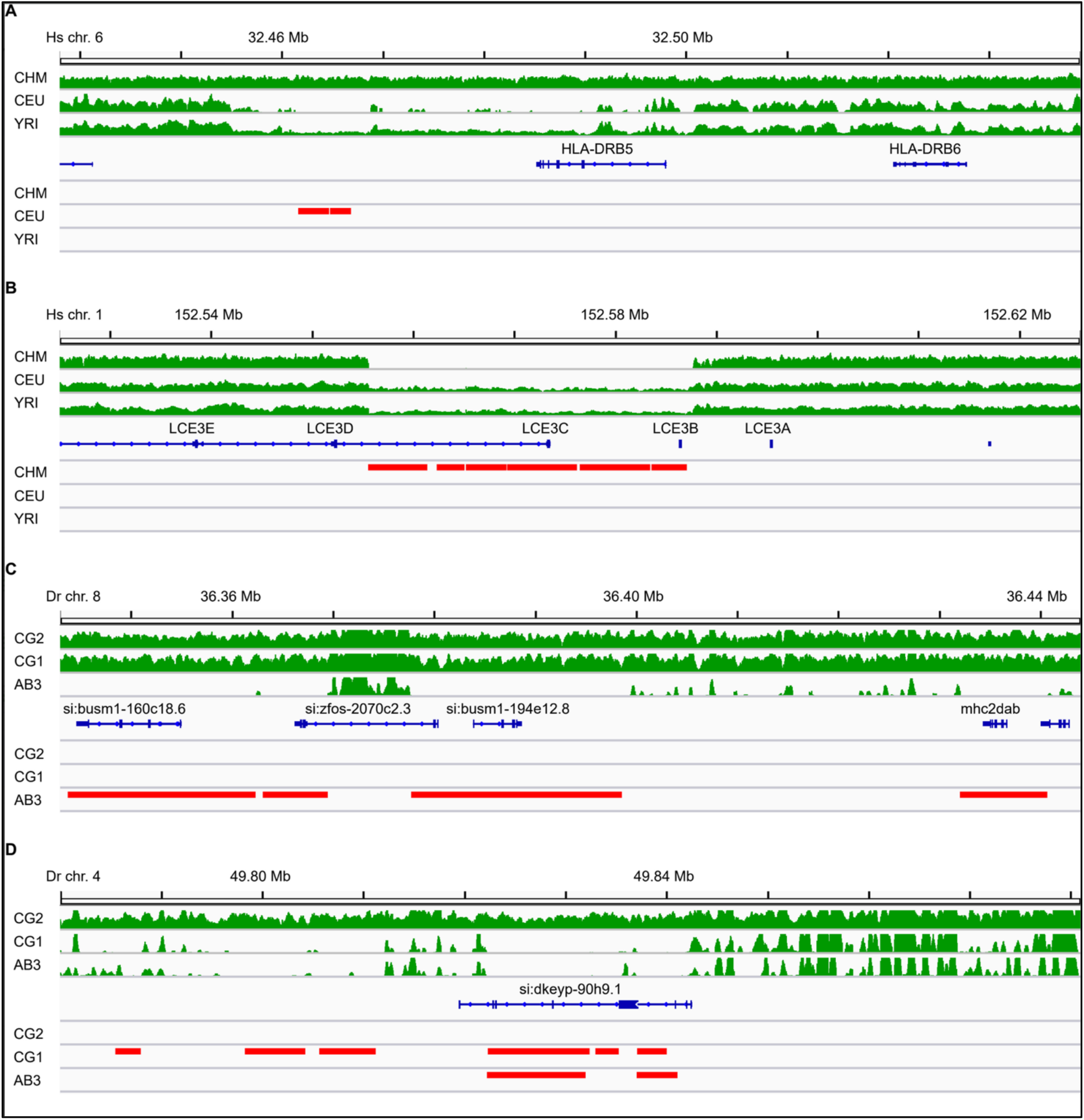
Zero Coverage Regions highlight unique haplotypes. Selected 100 kb region plots including (A) *HLA-DRB5* gene on human chromosome 6, (B) *LCE3C* gene on human chromosome 1, (C) *mhc2dab* gene on zebrafish chromosome 8, and (D) NLR gene (*si:dkeyp-90h9*.*1*) on zebrafish chromosome 4. Plots show mapped sequence read coverage across each region in green. Zero Coverage Regions (no mapped reads over ≥ 2 kb intervals) are displayed in red.

In contrast to human MHC genes, which appeared to lack direct overlap with ZCRs, many zebrafish MHC genes were found to overlap ZCRs, including the MHC Class II gene *mhc2dab* (Fig 4C). We hypothesized that ZCRs might highlight sequences that are either altogether missing, or instead highly divergent. To identify alternative haplotypes that might be associated with divergent sequences, we performed genomic assembly for the three zebrafish individuals using Discovar de novo. Analysis of these assemblies with BUSCO (Supplementary Table 5) returned high percentages of the target genes for each assembly, comparable to the zebrafish reference genome, particularly for the CG1 and CG2 assemblies (85-86% complete genes). Assembly metrics (Supplementary Table 6) indicated high quality assemblies including N50 values of 30-40 kb for the CG1 and CG2 assemblies and 16 kb for AB3. BLAST searches of these assemblies identified scaffolds either highly similar to the reference, or highly divergent, depending upon the sample.

Examining variation at the haplotype level, including alignment of scaffolds from our genomic assemblies, revealed that while reference or highly similar haplotypes were found across many loci, in other cases samples lacked reference haplotype sequences over large regions (Supplementary Figs 2-5). This was particularly evident for the zebrafish chromosome 4 region associated with NLR genes (Fig 4D), where a highly variable patchwork of different haplotypes was apparent and where often only one of the samples carried reference or similar haplotypes across an NLR gene cluster.

Our finding of a ZCR overlapping *mhc2dab* was somewhat unexpected, given that this gene has been considered the lone classical MHC Class II beta gene in zebrafish. However, we noticed a pattern in coverage throughout the larger MHC Class II locus where reads were missing over a large segment (∼100 kb including *mhc2dab* around 36.4 Mb) for the AB3 fish, and reads were present only in the AB3 fish for an even larger portion (∼200 kb including *mhc2dgb* around 35.3 Mb) that were missing in the other fish (Fig 5A). We hypothesized that these represent alternative haplotypes within the MHC Class II locus on zebrafish chromosome 8, which appear to be represented as a composite in the reference genome.

**Figure 5.**
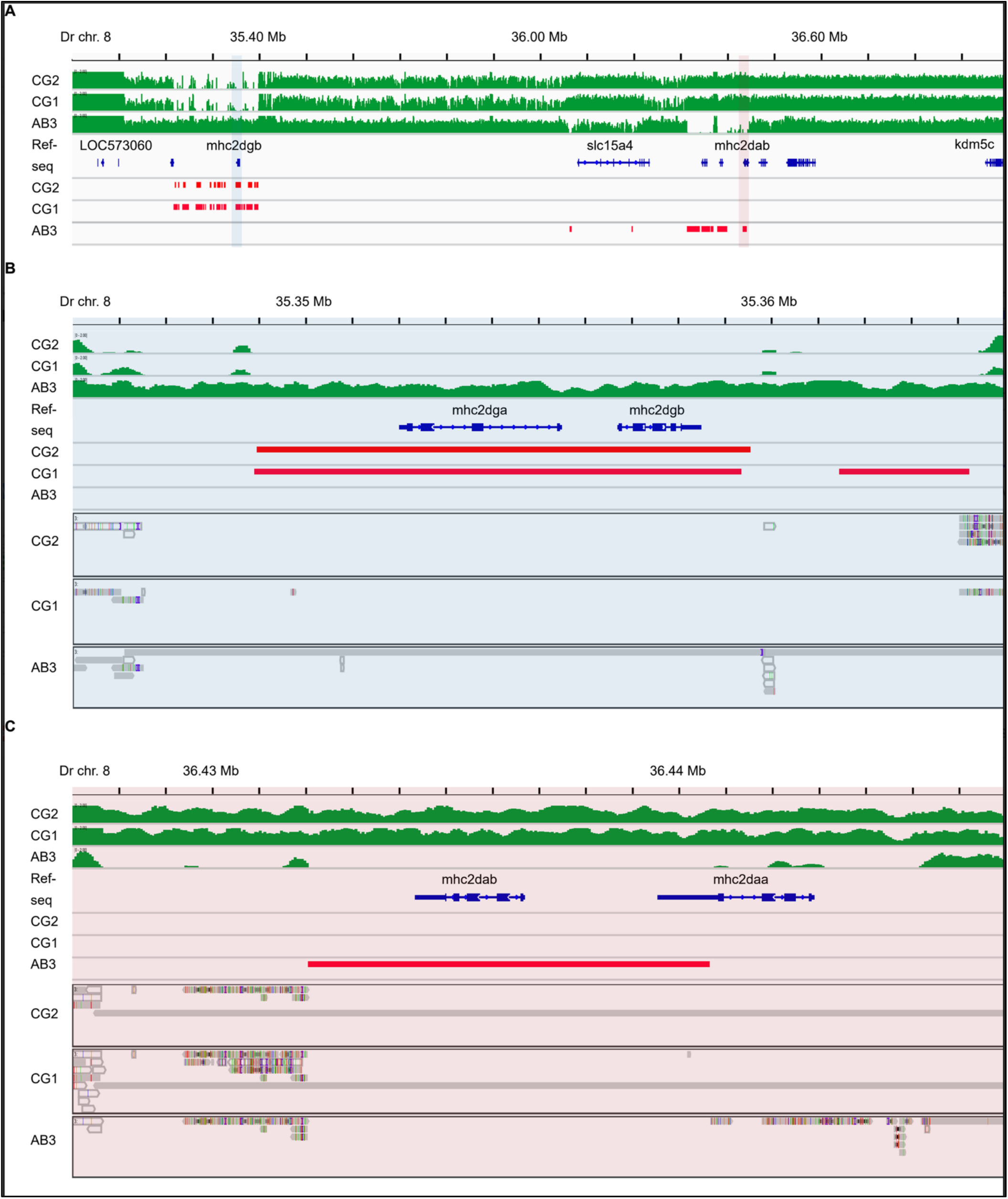
MHC Class II genes on zebrafish chromosome 8. (A) Read coverage across the zebrafish Class II MHC locus shows marked variability between individuals. Unlike the CG2 and CG1 fish, the AB3 zebrafish genome has a cluster of Zero Coverage Regions (ZCRs, without any mapped reads over ≥ 2 kb intervals) in the region surrounding *mhc2dab* (highlighted in light red). In contrast, the CG2 and CG1 fish have a cluster of ZCRs in the region surrounding *mhc2dgb* (highlighted in light blue). (B) A more detailed view of the region highlighted above in blue (A) showing ZCRs overlapping the neighboring *mhc2dgb* and *mhc2dga* genes. (C) A detailed view of the region highlighted above in red (A) showing ZCRs overlapping the neighboring *mhc2dab* and *mhc2daa* genes. Read coverage is depicted in green, ZCRs are in red, and scaffolds from Discovar assemblies that align to the reference genome are grey.

BLAST searches of our genomic assemblies returned scaffolds nearly identical to the reference sequence for *mhc2dab* for CG1 and CG2 (Supplementary Table 7). On the other hand, for AB3 the closest scaffold match contained *mhc2dgb*, a gene with high amino acid identity to *mhc2dab* (81%), and high expression levels (Supplementary Table 8). We had similar findings for *mhc2dga* in AB3, with high amino acid identity to *mhc2daa* (64%), and high expression, consistent with an MHC Class II classical gene signature.

Thus, only the AB3 fish had coverage data and genomic scaffolds consistent with reference sequence that encompassed *mhc2dga* and *mhc2dgb* (Fig 5B). On the other hand, only the CG1 and CG2 fish had coverage data and genomic scaffolds including *mhc2dab* and *mhc2daa* (Fig 5C). This pattern is reminiscent of our earlier observation of alternative haplotypes for the MHC Class I locus (McConnell et al. 2016), providing evidence in this case that two alternative MHC Class II haplotypes appear to be represented in tandem within the zebrafish reference genome.

Because pathway analysis implicated zebrafish immune genes as highly enriched among genes with evidence of positive selection and genes associated with ZCRs, we elected to more comprehensively examine the association with MHC and NLR genes. We used custom gene lists (Supplementary Tables 9-11) because these genes often lacked RefSeq annotation. Strikingly, 62% of the MHC gene set (Fig 6A) and 63% of the NLR gene set (Fig 6B) were associated with ZCRs in at least one of the three zebrafish samples, compared with 0% for MHC and NLR genes in humans. When taking all genes into consideration, 5% (n=1461) of zebrafish genes had exons overlapping with ZCRs (Fig 6C, Supplementary Tables 12-21), while less than 0.2% (n=36) of all human genes overlapped ZCRs in at least one sample. This high level of presence/absence variation in zebrafish individuals is expected to disproportionately affect immune function given the large number of immune genes involved.

**Figure 6.**
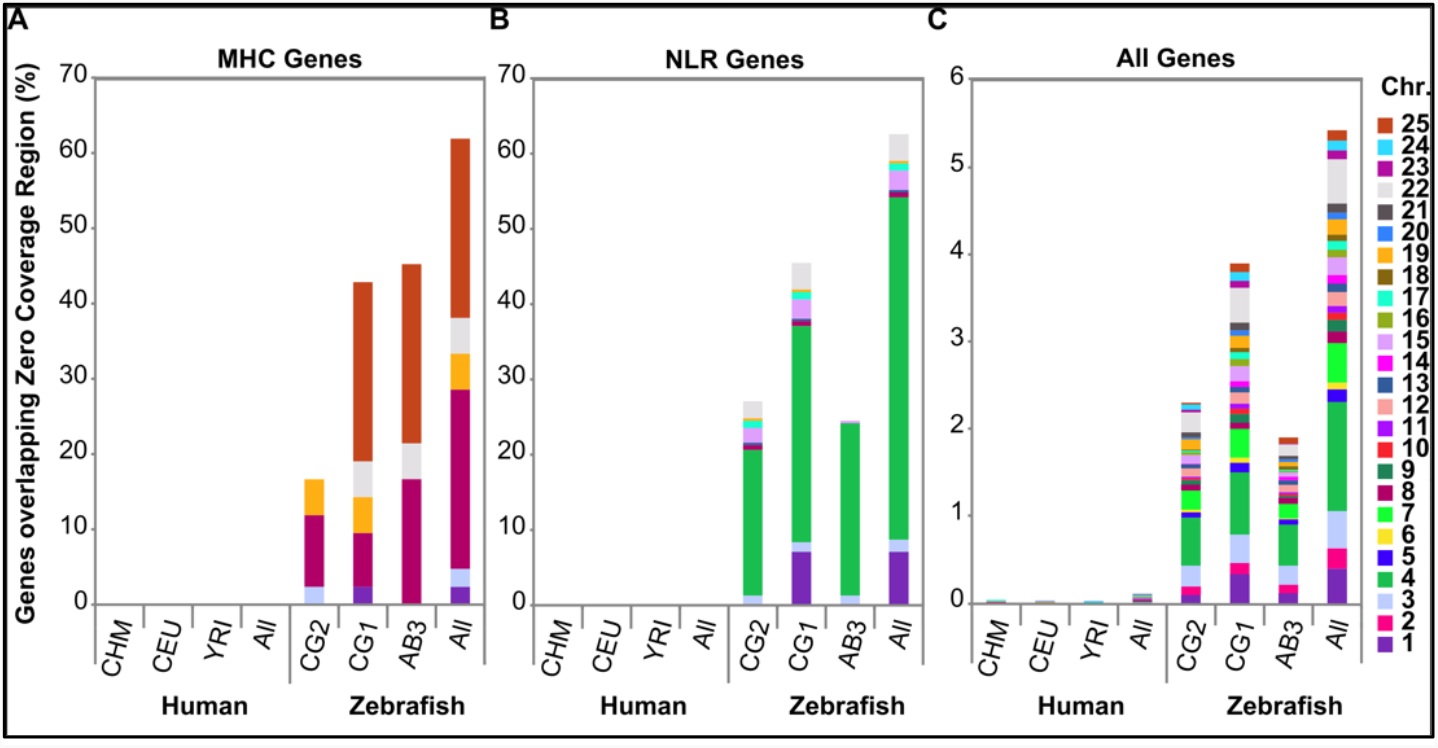
Immune genes associated with Zero Coverage Regions (ZCRs). The percentage of (A) Major Histocompatibility Complex (MHC) genes, (B) NOD-like receptor (NLR) genes, or (C) all genes in each of three human or zebrafish genomes, with at least one exon overlapping ZCRs. ‘All’ represents the cumulative number of unique genes for each type that overlaps with ZCRs across all three individuals. CHM, CEU, and YRI are samples from the 1000 genomes project representing: a haploid complete hydatidiform mole, CHM1; Utah Resident (CEPH) with European Ancestry, NA12878; Yoruba in Ibadan, Nigeria, 19240; respectively. CG2 and CG1 are clonal zebrafish lines, and AB3 is a partially inbred fish, all on the AB genetic background.

## DISCUSSION

We have found that zebrafish genomes carry much higher levels of variation than human genomes. This was evident even when comparing homozygous diploid clonal fish (effectively haploid) genomes to human genomes carrying relatively high levels of variation, for example the genome of an individual of African ancestry. However, our analysis likely represents an underestimate of variation due to challenges in characterizing divergent zebrafish gene loci, including under-sampling and inherent limitations of reference genomes.

Previous studies of zebrafish genetic diversity have revealed a high density of copy number variants (CNVs) and single nucleotide variants (SNVs) (Brown et al. 2012; Guryev et al. 2006). Unlike previous studies of zebrafish genomic variants which found deletions to be more prevalent than insertions (Patowary et al. 2013; LaFave et al. 2014), we observed no significant bias detecting insertions and deletions, leading to an observed insertion/deletion ratio that closely matches their expected (equal) distributions throughout the zebrafish genome. In our study, 99.4% of insertions were identified as novel, compared with only 62.9% of deletions (Supplementary Table 2), likely because of improved variant detection, particularly for insertions.

Due to challenges associated with using CNV-calling pipelines for zebrafish sequencing data, including higher sequence divergence and non-uniform patterns, we used ZCRs to highlight regions with no mapped reads as an unbiased method to identify putative deletional CNVs. Our findings suggest that while ZCRs were highly specific to identify large regions with missing coverage, they were not sensitive to detect all deletions and/or regional polymorphism in genes. This was particularly evident when read coverage was more variable due in part to mapping of highly related gene sequences, or when samples were heterozygous for such variation. ZCRs are thus overall likely to underestimate the degree of structural variation throughout these genomes, including alternative haplotypes which will likely require long-read sequencing or similar approaches to fully resolve.

This may be particularly relevant for immune genes, as gene pathway analysis showed that much of the zebrafish variation was concentrated in genes associated with immune function, whereas human variation was associated with other gene pathways. MHC genes, arguably the most polymorphic genes in humans, also exhibited variability in zebrafish genomes that far exceeded MHC variation found in humans (Fig 6A). Even nonclassical MHC genes, which are largely monomorphic in humans, were also found to be highly polymorphic in zebrafish, consistent with more widespread differences in immune genes between individuals.

On chromosome 4, which is highly enriched for NLR genes (Howe et al. 2016), we found that roughly half of the chromosome appeared largely different from one individual fish to the next. The magnitude of this variation was particularly striking for the CG1 fish, where reads across nearly half of chromosome 4 failed to align to the reference genome. Yet, most of these segments (e.g., ∼ 1 Mb blocks) still had high sequence similarity to the reference genome in one or more of the other fish, indicating that our alignment approach worked well for the sequences present in these other samples. Poorly mapped or missing reads, as outlined by ZCRs, were concentrated throughout these large segments of chromosome 4 that varied markedly between individuals. Our genomic assemblies for the individual zebrafish provide additional evidence that in many cases these poorly mapped reads were due to markedly divergent sequences (or alternatively, presence/absence variation) and not due to low quality sequence data. Despite annotation efforts to define the scope of the NLR genes in the reference genome (Howe et al. 2016), further work is needed to uncover additional genes in alternative haplotypes that we identified, each spanning up to 20 Mb. These genes were underreported in our assessment of positive selection, due to annotation being incomplete. Analysis of the genes in these ZCR regions using RNA-Seq data across different tissues would clarify expression patterns, provide insight into their function, and further improve annotation to include genes that may not be present in current reference genome sequences.

The functional implications of such expanded and diverse repertoires of NLR genes, along with any consequences for evolution of the host genome, remain interesting topics for further study. Species can gain distinctive collections of immune genes, which allow them to respond to the evolving threats of pathogens. Some immune genes have been found to segregate within only certain individuals within a species (Horton et al. 2008; McLure et al. 2013), including in zebrafish (McConnell et al. 2014). While previous studies of vertebrate NLR genes have focused on differences between species (Jones et al. 2016; Tørresen et al. 2018), here we find that the proliferation of zebrafish NLR genes appears highly variable between individuals. Intriguingly, plant NLRs also maintain strain-specific complements of NLR genes, which are known to help mediate strain-specific pathogen resistance (Maekawa et al. 2011; Meunier and Broz 2017). While the functional roles of these highly variable zebrafish NLR gene sets remain unclear, they may be analogous to earlier findings from plants and other non-vertebrate species.

To our knowledge this scale of immune gene diversity between individuals has not previously been described in vertebrates. Thus, this study adds to our understanding of immune gene evolution by highlighting divergent sequences comprising half of a vertebrate chromosome, where hundreds of genes may vary markedly between individuals. Future studies are needed to examine how this variation may impact immune gene function in this vertebrate model, offering a unique opportunity to study ongoing mechanisms driving large-scale genome evolution.

## METHODS

### Zebrafish

The golden-derived clonal lines CG1 (Smith et al. 2010) and CG2 (Mizgirev and Revskoy 2010), were each generated through two rounds of parthenogenesis and generously provided by Dr. Sergei Revskoy. The AB3 individual zebrafish from the AB zebrafish strain was also kindly provided by Dr. Sergei Revskoy. Zebrafish husbandry, care and all experiments were performed as approved by the University of Chicago Institutional Animal Care and Use Committee.

### Genomic sequencing

We used a single-library per sample approach for high-throughput sequencing. Briefly, Illumina TruSeq DNA PCR-free libraries were constructed from genomic DNA isolated from individual zebrafish. To facilitate Discovar *de novo* assemblies, the libraries were individually sequenced in single lanes on a HiSeq2500 instrument (Rapid run mode), using paired-end 2 × 250 bp reads, providing approximately 50-60x coverage.

### Read Alignment

Zebrafish raw reads were aligned to the GRCz10 assembly (Illumina iGenomes: https://support.illumina.com/sequencing/sequencing_software/igenome.html) using BWA aln v0.7.12 with aln parameters: *-q 5 –l 32 –k 2 –o 1*; sample parameters: *-a 1350* (Li and Durbin 2009) and formatted using sambamba v0.5.9 (Tarasov et al. 2015).

### SNV/indel Detection

D. rerio alignments were filtered to remove unaligned reads and alignments with low mapping quality (MAPQ > 10) using sambamba. Filtered alignments were base quality recalibrated using GATK v3.6.0 (McKenna et al. 2010). Filtered and quality recalibrated alignments were used to detect genotypes using the GATK HaplotypeCaller and GenotypeGVCFs tools. To call genotypes, haplotypes were first detected in each sample separately then joint-genotyping was performed across all three samples using the GATK HaplotypeCaller/GenotypeGVCFs. Raw genotypes were hard filtered to remove low quality calls and potential artifacts using GATK’s SelectVariants and VariantFiltration (Supplemental Table 1). Basic variant metrics were extracted using RTG Tools v3.7.1 (Cleary et al. 2014, 2015) and custom scripts. Filtered variants were annotated using the ENSEMBL’s VariantEffectPredictor (VEP) v85 (McLaren et al. 2016) with RefSeq cache version 85.

### dN/dS Analysis

VEP annotations were processed to select the mutational impact on the canonical transcript for each alternate allele. Synonymous and non-synonymous effects were then counted for each gene based on the canonical transcript and imported into R. The ratio of non-synonymous to synonymous counts (dN/dS) for each gene was estimated. Genes with dN/dS > 1.0 were used for an enrichment analyses with the clusterProfiler v3.2.15 (Yu et al. 2012) and DOSE v3.0.10 (Yu et al. 2015) R BioconductoR packages.

### Human Sample Coverage and Variant Data

The human alignment files were downloaded from the 1000 Genomes FTP site:

/1000genomes/ftp/phase3/data/NA12878/high_coverage_alignment/NA12878.mapped.ILLUMIN

A.bwa.CEU.high_coverage_pcr_free.20130906.bam

/1000genomes/ftp/phase3/data/NA19240/high_coverage_alignment/NA19240.mapped.ILLUMIN

A.bwa.YRI.high_coverage_pcr_free.20130924.bam

/vol1/ftp/technical/working/20150612_chm1_data/alignment/150140.mapped.ILLUMINA.bwa.CH

M1.20131218.bam

The human VCF files were downloaded from: ftp://ftp-trace.ncbi.nih.gov/1000genomes/ftp/technical/working/20140625_high_coverage_trios_broad/ ftp://hengli-data:lh3data@ftp.broadinstitute.org/hapdip/vcf-flt/CHM1.mem.hc-3.3.flt.vcf.gz (The 1000 Genomes Project Consortium, 2005)

### Zero Coverage Region (ZCR) Analysis

Unfiltered BAM files were converted to 1x-coverage bigWig files using deeptools v2.4.3 (Ramírez et al. 2014). Gap regions were extracted from the UCSC table browser and removed from the bigWig files using bwtool v1.0-gamma (Pohl and Beato 2014). Regions from gap-removed bigWig files with 0 coverage were extracted and converted to BED files using bwtool and those >=2kb in length were extracted for downstream analysis. The selected regions were intersected with GTF files and the genes with at least one exon overlapping were extracted using the pybedtools v0.7.9 python package (Dale et al. 2011; Quinlan and Hall 2010) and custom scripts. Genes with overlapping ZCRs were then used for enrichment analyses in a similar manner as the dN/dS analysis.

### Genomic Assemblies Generated using Discovar de novo

Raw reads were converted to the unmapped bam format using Picard tools (2.2.1; http://broadinstitute.github.io/picard/). Discovar de novo (Weisenfeld et al. 2014) was used to generate genomic assemblies with default settings (build 52488; https://www.broadinstitute.org/software/discovar/blog/). While the Discovar de novo assemblies were each generated independently of the reference genome, the GRCz10 zebrafish assembly (version 140) was subsequently referenced for the purposes of scaffold mapping.

### BUSCO assembly metrics

Discovar de novo assemblies were analyzed using BUSCO (Benchmarking Universal Single-Copy Orthologs) (build v1.22 depending on Augustus v 3.1, blast+2.2.31, and hmmer3.1b2; http://busco.ezlab.org/), modified to run tblastn outside of the BUSCO script. The BUSCO approach provides quantitative assessment of genome quality by assessing genome completeness, based on an evolutionarily conserved list of 3023 vertebrate single-copy orthologs (Simão et al. 2015). Because we found that the tBLASTn results were sometimes incomplete using the implementation provided by the BUSCO genome assemblies, we instead performed our own tBLASTn searches on our genome assemblies using a separate installation. Complete tBLASTn results for each of our genome assemblies were then returned to BUSCO for gene prediction and assessment of completeness. We also included the GRCz10 reference genome in this modified BUSCO pipeline for comparison.

## DATA ACCESS

Genomic assembly data generated in this study have been submitted to the NCBI BioProject database (https://www.ncbi.nlm.nih.gov/bioproject/) under accession numbers PRJNA292113, LKPD02000000 (CG2); PRJNA454110, JALCZS000000000 (CG1); and PRJNA454111, JALCZT000000000 (AB3). Raw sequence data have been deposited in the NCBI short read archives (SRA) with accession numbers SRR7080552, SRR7081528, and SRR7081557. Supplemental data files have been published in the CyVerse Data Commons under https://de.cyverse.org/data/ds/iplant/home/shared/commons_repo/curated/McConnell_ZeroCoverageRegions_2022.

## ACKNOWLEDGEMENTS

The authors would like to thank Wilfredo Marin for excellent fish care and technical support and Sergey Revskoy for generously sharing zebrafish. We also thank Pieter Faber from the University of Chicago Genomics core for sequencing expertise and Cancer Center Support Grant (P30 CA014599) for sequencing support. This work was funded, in part, by a Postdoctoral Research Grant from the Chicago Biomedical Consortium (S.C.M.), with support from the Searle Funds at the Chicago Community Trust (J.L.O.d.), and the University of Chicago Cancer Research Foundation Auxiliary Board (J.L.O.d.).

